# Social Stress Alters Immune Response and Results in Higher Viral Load During Acute SIV Infection in a Pigtailed Macaque Model of HIV

**DOI:** 10.1101/2020.04.21.054130

**Authors:** Selena M. Guerrero-Martin, Leah H. Rubin, Kirsten M. McGee, Erin N. Shirk, Suzanne E. Queen, Ming Li, Brandon Bullock, Bess W. Carlson, Robert J. Adams, Lucio Gama, David R. Graham, Christine Zink, Janice E. Clements, Joseph L. Mankowski, Kelly A. Metcalf Pate

## Abstract

While social distancing is a key public health response during viral pandemics, social stress, which can be induced by social isolation, has been implicated in adverse health outcomes in general^1^ and in the context of infectious disease, such as HIV^2,3^. A comprehensive understanding of the direct pathophysiologic effects of social stress on viral pathogenesis is needed to provide strategic and comprehensive care to patients with viral infection. To determine the effect of social stress on HIV pathogenesis during acute viral infection without sociobehavioral confounders inherent in human cohorts, we compared commonly measured parameters of HIV progression between singly and socially housed SIV-infected pigtailed macaques (*Macaca nemestrina*). Singly housed macaques had a higher viral load in the plasma and cerebrospinal fluid and demonstrated greater CD4 T cell declines and greater CD4 and CD8 T cell activation compared to socially housed macaques throughout acute infection. These data demonstrate that social stress directly impacts the pathogenesis of acute HIV infection and imply that social stress may act as an integral variable in the progression of HIV infection and potentially of other viral infections.

## Main Text

Social stress is an environmental factor that has a measurable impact on general health outcomes^1^. Today, social stress has taken on greater societal significance, with social distancing practices vital to slow the SARS-CoV-2 pandemic^4–6^. The logical next question of how social stress impacts the body’s responses to pathogens, such as viral infections, remains incompletely understood. While data is currently unavailable for analysis in the COVID-19 pandemic, other viral pandemics, such as human immunodeficiency virus (HIV), may provide insight. HIV infection remains widespread and outcomes for PWH vary greatly and depend upon a number of factors, including social stress. PWH are at high risk for social stress due to stigma associated with infection and marginalization of people from demographics that are considered at higher risk of infection^7^. Not surprisingly, PWH report decreased physical and mental well-being compared to the general population^2^, and worse mental and physical quality of life than individuals with other chronic illnesses^3^.

Determination of the direct physiologic effects of social stress on viral pathogenesis and mechanisms through which such stress affects outcomes are complicated by the multifaceted nature of social stress and associated sociobehavioral confounders that have the potential to limit access to effective care or otherwise compromise health. The present study aims to determine if social stress directly affects HIV pathogenesis by comparing how single housing affects the progression of acute SIV infection in pigtailed macaques (*Macaca nemestrina*) in a retrospective analysis. Simian immunodeficiency virus (SIV) infection in macaques recapitulates many aspects of HIV pathogenesis, including decline in circulating CD4 T cells following infection and persistence of virus in latent reservoirs despite ART^8^. We hypothesized acute SIV infection would have greater impact on macaques housed individually prior to and following viral inoculation compared to macaques socially housed in pairs or trios. We demonstrate that housing macaques individually directly affects SIV viral load, CD4 T cell decline and T cell activation during acute infection. Social stress thus directly influences disease progression in the SIV-infected macaque model of HIV, and social stress is likely to have direct pathophysiologic impact on health outcomes of PWH and potentially on the pathogenesis of other viral infections.

## Results

### Social Stress Results in Elevated Viral Loads and Greater CD4+ T Cell Decline During Acute SIV Infection

Viral load and CD4+ T cell decline are two of the most frequent parameters used to assess clinical progression in people with HIV. To determine if social stress directly influences the progression of HIV, we used linear mixed effects regression models to conduct a retrospective analysis of data generated throughout acute infection (days 7, 10 and 14 postinoculation) by singly housed (n=35) compared to socially housed (n=41) macaques dual-inoculated intravenously with SIV/DeltaB670 and SIV/17E-Fr. This protocol of SIV infection has been shown to lead to high levels of viremia in the plasma and cerebral spinal fluid (CSF) and progression to AIDS defining criteria in 3 months^9^. We assessed the SIV viral load in the plasma and CSF using qRT-PCR to detect SIV gag. Singly housed macaques had higher plasma viral loads than socially housed counterparts throughout acute infection (by 6.1%, 6.5% and 21% on days 7, 10 and 14, respectively) (Fig.1A; p<0.0001). Singly housed macaques also had higher viral loads in the CSF than socially housed macaques (13%, 12% and 18% difference at the same timepoints) (Fig.1B; p<0.0001). The magnitude of the difference in viral loads between the singly and socially housed macaques compounded over time in the plasma (p=0.0007) but not the CSF.

**Figure 1.**
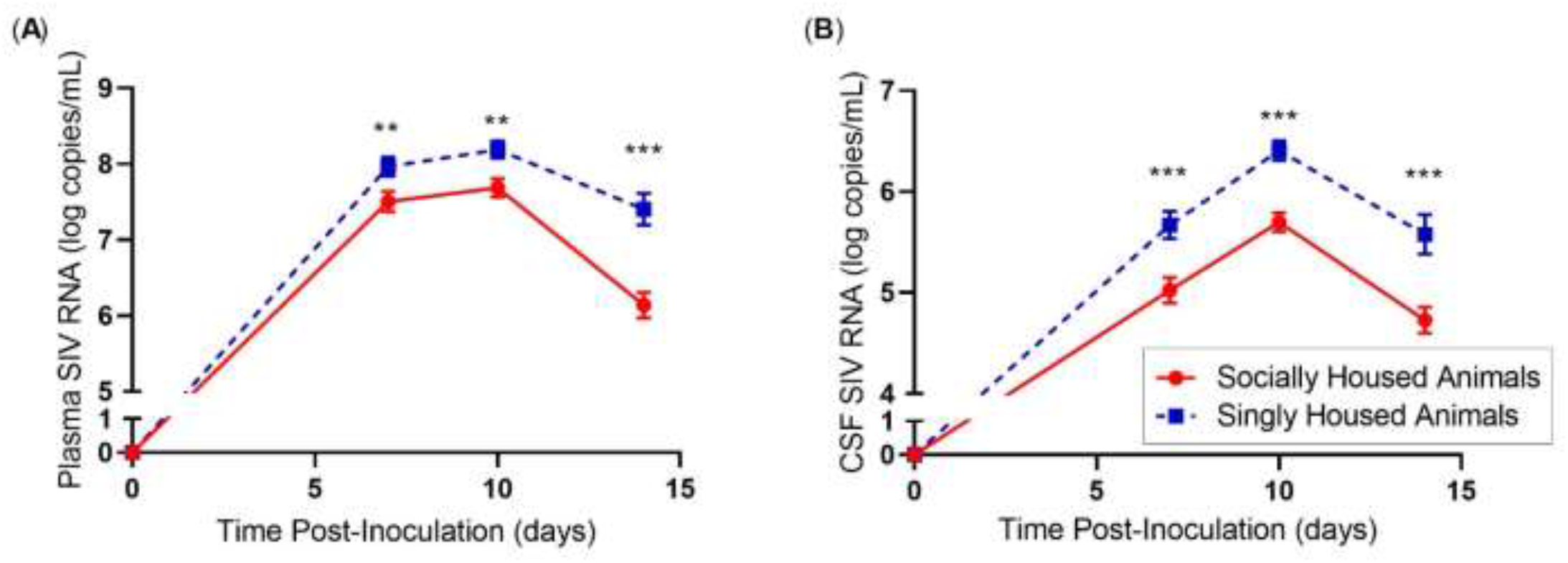
Social stress increases the plasma and CSF viral load in acute SIV-infection. SIV viral load in plasma (A) and cerebrospinal fluid (CSF) (B) were quantified utilizing qRT-PCR. Linear mixed effects regression model. Statistics depicted in figure represent significance of difference between singly and socially housed animals at each time point. \**P*≤0.05, \*\**P*≤0.01 and \*\*\**P*≤0.001. Circles connected by a solid line (red) indicate socially housed animals (N=41) while squares connected by a dashed line (blue) represent singly housed animals (N=35). Error bars represent the standard error of the least square mean estimates generated by the linear mixed effects regression model. ART initiated on day 12 post-inoculation.

We used flow cytometry coupled with complete blood counts to examine absolute T cell numbers in these same animals. As expected, all macaques demonstrated a decline in CD4 T cell numbers upon infection (Fig. 2A; p<0.0001). Singly housed pigtailed macaques had a 38.5% greater decline in CD4 T cells following SIV inoculation compared to socially housed animals (Fig. 2A; p=0.0001). Additionally, singly housed animals had a 34.1% greater decline in CD8 T cells (Fig. 2B; p=0.0002) and a 44.4% greater decline in total circulating lymphocytes (Fig.S1A; p<0.0001). These differential declines originated pre-inoculation, during which CD4 T cell numbers were 27.8% higher (Fig.2A; p=0.007), CD8 T cells were 25.9% higher (Fig.2B, p=0.0006) and total lymphocytes were 28% higher (Fig.S1A; p=0.0004) in singly housed compared to socially housed animals. A lower CD4 to CD8 T cell ratio is associated with negative outcomes in PWH10, and, as expected, infection led to lower CD4/CD8 ratios in all animals (p=0.0004). However, the CD4/CD8 ratio was 18% lower in singly housed animals at day 14 following inoculation compared to socially housed animals (Fig. S1B; p=0.04).

**Figure 2.**
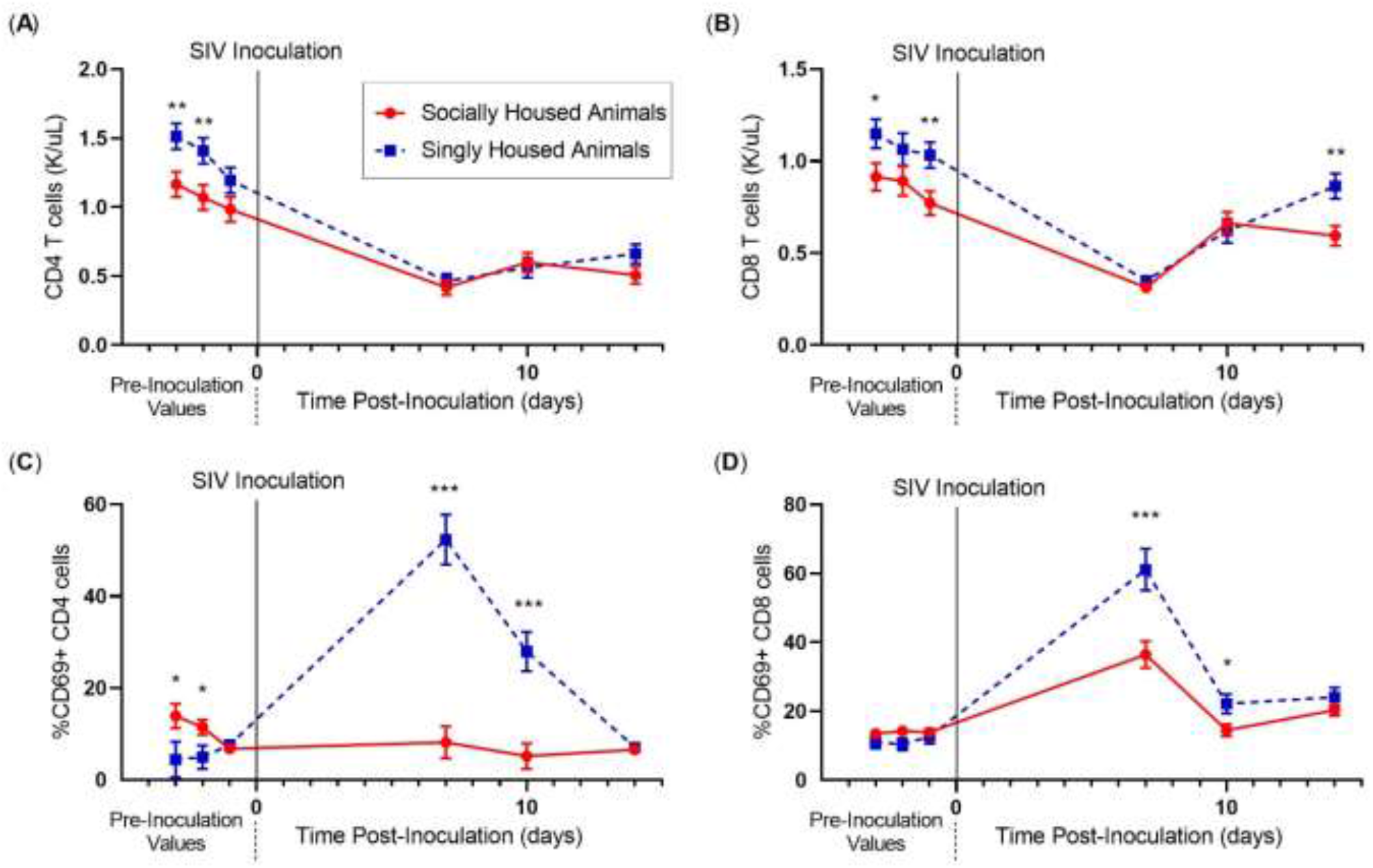
Social stress increases the immune impact of acute SIV-infection. CD4 T cells (A), CD8 T cells (B) in the peripheral blood of macaques pre- and post-inoculation with SIV. Percentage of CD4 T cells expressing CD69 (C) and percentage of CD8 T cells expressing CD69 (D). Linear mixed effects regression model. Statistics depicted in figure represent significance of difference between singly and socially housed animals at each time point. \**P*≤0.05, \*\**P*≤0.01 and \*\*\**P*≤0.001. Circles connected by a solid line (red) indicate socially housed animals (N=41) while squares connected by a dashed line (blue) represent singly housed animals (N=35). Solid vertical line at day 0 indicates time point of SIV-inoculation. Error bars represent the standard error of the least square mean estimates generated by the linear mixed effects regression model. ART initiated on day 12 post-inoculation.

### Social Stress Alters the Immune Response During Acute SIV Infection

To further determine the effect of social stress on the immune response to SIV infection, we examined CD4 and CD8 T cell activation. Activated CD4 T cells are susceptible to infection with HIV and are key producers of virus during acute infection^11,12^. Notably, singly housed macaques had 9.2-fold more CD69+ CD4 T cells at peak activation in acute infection compared to pre-inoculation values (Fig.2C; p<0.0001) whereas socially housed animals demonstrated negligible activation. CD8 T cells are essential for viral control, and CD8 T cell activation is an expected sequelae of acute HIV infection^13,14^. All macaques had heightened CD8 T cell activation during acute infection, demonstrated by increased percentage of cells expressing CD69, with peak activation on day 7 (Fig.2D; p=0.001). However, singly housed animals had 68% more CD69+ CD8 T cells at peak activation compared to socially housed (p=0.0007).

### Social Stress Increases the Variability of the Data Produced by SIV-Infected Macaques

Reduction of exogenous stress through the introduction of other stress-reducing refinements has been shown to reduce the variability of data in macaque models15. To evaluate if social stress similarly impacts data variability in this SIV-infected pigtailed macaque model, we compared the standard deviations of all the post-inoculation data reported herein between socially and singly housed animals. Data produced by singly housed animals was overall more variable than the data from socially housed animals (Fig.3 and S2; p=0.02).

**Figure 3.**
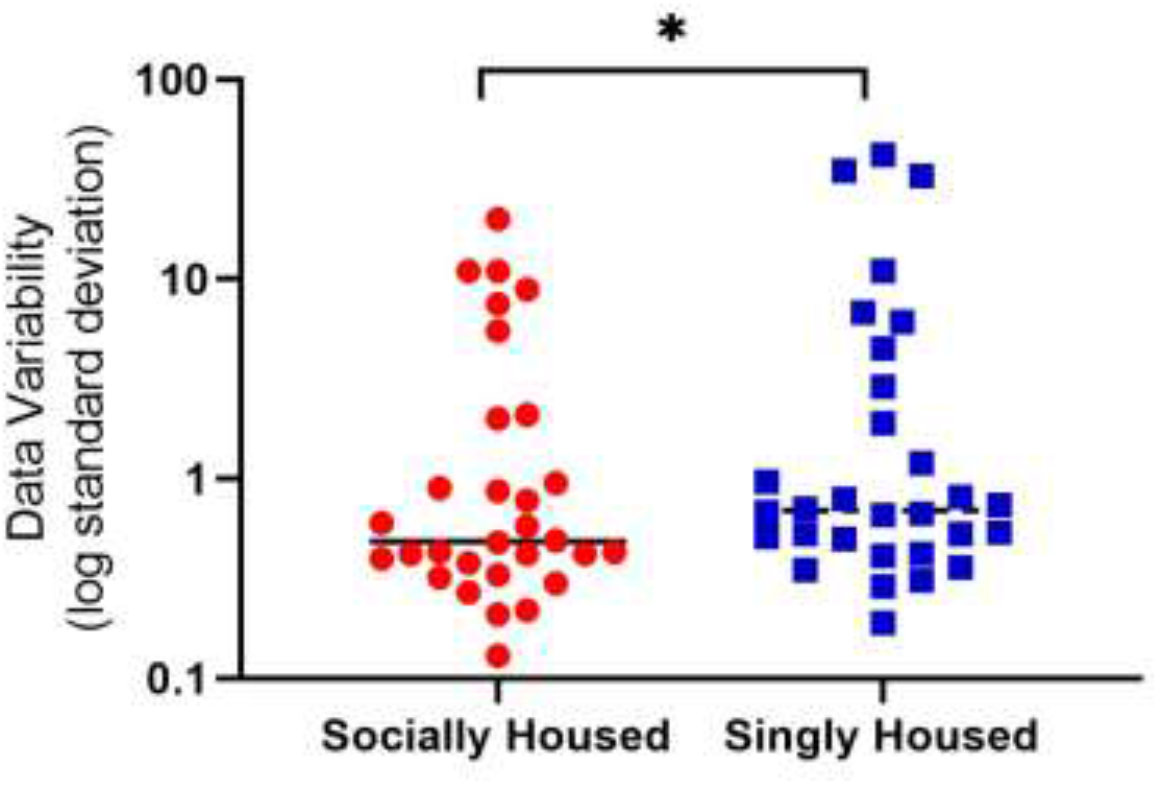
Social stress increases the variability of data produced by the SIV-infected macaque model during acute infection. Standard deviation for all data sets presented in Figures 1, 2, and S1 were compared in a pair-wise fashion between socially (N = 30 parameters for 41 animals) and singly (N = 30 parameters for 35 animals) housed animals using a twotailed Wilcoxon matched pairs signed rank test. \**p*≤0.03.

## Discussion

Our data demonstrate that reducing social stress by providing social housing with a compatible conspecific lessens the immune impact of acute SIV infection in a pigtailed macaque model of HIV infection, resulting in lower plasma and central nervous system (CNS) viral loads, a less marked decline in circulating CD4 T cells and a reduction in CD4 and CD8 T cell activation. This effect is observed at the earliest timepoints examined and throughout acute infection while all other aspects of infection and care remained consistent between these groups (Table S1), effectively removing the sociobehavioral confounders that affect access to care and adherence to treatment inherent in PWH. Thus, this study demonstrates that there is a direct pathophysiologic effect of social stress on the pathogenesis of HIV infection and implies that social stress may have implications for the pathogenesis of other viral infections as well.

This analysis has implications beyond primary infection for PWH. Latent reservoirs are established during acute infection, remain the major barrier to cure, and are especially difficult to eliminate from sanctuary sites such as the brain^16,17^. Our findings of higher viral loads in the plasma and CSF in the context of social stress have potential long-term implications for PWH. These higher viral loads may ultimately originate from the increased CD4 activation that we observed in the singly housed animals, as activated CD4 T cells simultaneously serve as permissive cells for the virus to infect and produce more virus if infected, thus potentiating viral spread and reservoir seeding^12^. Further, the CD4/CD8 ratio was lower late in acute infection and CD4 decline greater in singly housed compared to socially housed animals. Lower CD4/CD8 ratio is a predictor of non-AIDS related morbidity and mortality^10^ and age-associated disease in PWH^18^, while low CD4 counts are a direct prognostic indicator of disease progression throughout infection^19^. Our observation of these negative prognostic indicators in the context of social stress provides additional evidence that social stress directly contributes to negative outcomes. Thus, this work provides important evidence that social stress may have a direct adverse effect on the immune response to HIV, viral production, the establishment of latent reservoirs, viral load, and disease outcomes.

Due to the retrospective nature of our analysis we did not evaluate the mechanism underlying the effect of social stress on SIV/HIV pathogenesis yet work on the biology of stress responses of people with and without HIV provides insight. In uninfected people, chronic stress dysregulates immune function^20^, leading to a suppression in antibody production^21^, a suppression in leukocyte proliferation^22^, and a reduction in virus-specific T cell and NK cell activity^23^. Social isolation is associated with under-expression of proinflammatory genes, including those involved in innate antiviral resistance through type I interferon responses^24^, and could directly affect immune response and viral control. Compounding these effects, PWH have generally been shown to have increased levels of plasma cortisol compared to uninfected controls^25–26^, and inflammatory cytokine interleukin-1 and HIV gp120 are implicated in mediating this increased level of cortisol by increasing levels of corticotropin-releasing hormone^27^. Clinical studies demonstrate that glucocorticoid receptors of PWH have decreased affinity for cortisol, resulting in glucocorticoid insufficiency despite increased circulating cortisol. Viral suppression with ART does not fully attenuate the dysfunction in the physiologic stress response in PWH, illustrating the complexity of the interaction between HIV and the stress response^28^. Interestingly, social support mitigates the immunosuppressive impact of glucocorticoid release in uninfected individuals^29^. However, the findings from studies on the effect of social support on PWH have been equivocal, and any impact of social stress on disease progression and health outcomes in PWH is compounded by the fact that social isolation is more prevalent in PWH compared to uninfected controls^30^. In sum, social stressors affect immune functioning and thus HIV progression, and robust, controlled studies are needed to better understand the pathophysiologic mechanisms underpinning these effects on PWH.

Though this study was retrospective, we were able to control for the majority of variables that could have affected our study outcomes (Table S1). The present analysis focused on the beginning of acute SIV infection in order to eliminate the ART regimen variability as a confounding factor. All macaques were untreated until day 12 post-inoculation, when the majority of macaques were treated with antiretroviral (ART) regimens that differed between groups because these macaques were a part of 18 different studies. The differences presented here between the singly and socially housed groups are not driven by the post-ART datapoint, and the day 14 post-inoculation data includes only the animals that initiated ART at day 12 post-inoculation to exclude the potential confounder of lack of treatment. Other variables that were not controlled for in our study design were controlled for in multivariate post-hoc statistical analyses and found to not influence the conclusions. Importantly to note, there are limitations to the extent that these data can be taken as a representative model for the general population, as only data from juvenile male pigtailed macaques were available. For example, there is substantial literature documenting sex differences in the immune and stress responses and in HIV-associated disorders, such as neurocognitive impairment^31^.

These data also have implications for the translational value and reproducibility of HIV research with macaque models. Macaque models are instrumental in achieving scientific advancement for treatment of PWH and continue to be at the forefront of discovery in HIV and in other infectious virus research^32–40^. However, an animal model’s value is only as great as the translational value of its data, and how we keep animals in research settings can impact translational value and the reproducibility of generated data^41^. Specifically, excessive physiologic stress as a result of housing conditions such as single housing has the potential to reduce the translational value of animal models. Single housing of mouse and rat models of epilepsy led to an increase in corticosterone and adrenocorticotropic hormone levels, a decrease in brain-derived neurotrophic factor levels and exacerbated the epilepsy phenotype including seizure frequency^42,43^, leading to the conclusion that social isolation of social species adds an additional variable that can alter the interpretation of the data. Macaques are social animals, and housing macaques individually similarly results in physiologic stress. Otherwise healthy macaques relocated from social to single housing show modulated immune responses for weeks to months thereafter, as demonstrated by lower CD4/CD8 ratios and less robust proinflammatory cytokine responses^44^. Such changes to the immune system driven by social stress have the potential to confound infectious disease studies. Certain psychosocial variables such as social stability lead to an increased risk of mortality from SIV infection^45^. Our data complements the existing literature by demonstrating differences in the immune response and viral loads in singly compared to socially housed SIV-infected macaques. Though we cannot directly demonstrate improvements in translation and reproducibility through this work, the magnitude of the decline in CD4 T cell counts that we observed in socially housed macaques during acute infection (two-fold) is more analogous to that seen in humans during primary HIV infection^46^ than the three-fold change observed in singly housed macaques. Further, the decrease in variability of the data generated by our socially housed macaques bodes favorably for study reproducibility. Social housing of macaques is therefore an important refinement that may improve the translational value and reproducibility of data obtained from macaque models of HIV infection, and caution must be taken when comparing the data from studies completed in singly and socially housed macaques.

In summary, we have demonstrated that social stress in the form of single housing of macaques has a direct negative impact on the pathogenesis of acute SIV infection, leading to elevated peripheral and CNS viral loads, a more pronounced decline in circulating CD4 T cells, a lower CD4/CD8 ratio, and an increase in T cell activation. These data indicate that social support for PWH may have a direct positive impact on disease progression beyond the positive psychobehavioral effects and practical benefits, and provides evidence supporting a direct pathophysiologic effect of this social support on outcomes for PWH. Further research is needed to define the implications of social isolation on the pathogenesis of other viral infections. These data should furthermore be considered as evidence for the direct effect that social housing has on this vital animal model of HIV infection, as refinements such as social housing have the potential to improve the reproducibility and translational value of biomedical research conducted with animal models.

## Materials and Methods

### Animals

76 juvenile male pigtailed macaques (*Macaca nemestrina*) were inoculated intravenously with the same stock of simian immunodeficiency virus (SIV) inoculum containing the neurovirulent clone SIV/17E-Fr and the immunosuppressive swarm SIV/DeltaB670, as previously described^9^. A summary of the covariants controlled for in this analysis can be found in Table S1. All macaques were seronegative for SIV, simian T-cell leukemia virus, and simian type D retrovirus prior to study. All macaques tested negative for the MHC class I allele, *Mane-A1*08401*. Macaques were sedated intramuscularly with 10 mg/kg ketamine at 3 pre-inoculation timepoints that occurred at least two weeks apart and on days 7, 10 and 14 post-inoculation to facilitate blood and cerebral spinal fluid (CSF) collection. On day 12, the majority of macaques started on antiretroviral therapy (Table S2); any animals that did not begin an ART intervention were excluded from the day 14 post-inoculation analysis. The data for each parameter at each time point was generated based on grouping a single, distinct sampling of each animal within the socially or singly housed group. All animals consumed commercial macaque chow (Purina 5038) and water ad libitum throughout the study, and all macaques were provided daily enrichment by a behaviorist. 35 macaques were singly housed upon assignment to study. These animals remained separated from direct physical contact with conspecifics prior to inoculation (approximately 2-month duration), and throughout the course of the study. All singly housed macaques were able to see, hear and smell conspecifics in the same room though they did not share a cage with a conspecific. 41 macaques were socially housed (placed in compatible pairs or trios) upon study assignment for at least 2-months prior to inoculation, including all pre-inoculation time points, and remained with their conspecific(s) after inoculation and throughout the study period. Grouping of individuals was completed under oversight of an animal behaviorist to ensure social compatibility and reevaluated periodically throughout the study; no socially housed animals showed signs of incompatibility or had to be separated during the course of collecting these data. These data represent a retrospective analysis of a total of 18 studies conducted over a 10-year period, with the 9 studies involving the singly housed macaques occurring during the first 5-year period, and the 9 studies involving the socially housed macaques occurring during the second 5-year period (Table S2). All procedures were approved by the Johns Hopkins University Institutional Animal Care and Use Committee and were conducted in accordance with guidelines set forth in the Animal Welfare Regulations and the Guide for the Care and Use of Laboratory Animals. These studies were all conducted within a fully AAALAC accredited facility.

### Quantification of SIV Viral

Load Viral RNA was isolated from plasma and cerebral spinal fluid (CSF) using the QuantiTech Virus Kit (Qiagen). Viral loads were quantified by quantitative reverse transcriptase polymerase chain reaction (qRT-PCR) using primers in the SIV gag region: forward 5’-GTCTGCGTCATCTGGTGCATTC-3’; reverse 5’-CACTAGGTGTCTCTGCACTATCTGTTTTG-3’; 5’-FAM/3’-Black hole-labeled probe 5’-CTTCCTCAGTGTGTTTCACTTTCTCTTCTG-3’. All viral RNA extractions and PCR reactions over the 10-year period were prepared by one specific, skilled technician within the laboratory.

### Complete Blood Count and Flow Cytometry

Citrated whole blood samples were used for complete blood count (CBC), which were completed on site using a Hemavet (Drew Scientific) hematology analyzer calibrated for use with pigtailed macaque samples, or a Procyte Dx Hematology Analyzer (IDEXX Laboratories) (Table S2). Samples were excluded from this analysis at particular time points if clotting was noted upon CBC analysis, thus the n for each parameter at each time point varied. To assess the lymphocyte composition of the macaque whole blood, samples were stained with fluorochrome-coupled CD3, CD4, CD8a and CD69 antibodies for 20 minutes at room temperature and fixed for 10 minutes with BD FACS Lysing Solution (Becton Dickinson, Franklin Lakes, NJ). The samples from all of the socially housed animals and 17 of the singly housed animals were stained using the antibodies listed in panel A of Table S3; the remaining 18 singly housed animals were stained using the antibodies in panel B of Table S3. The stained whole blood samples were analyzed by a single specific, skilled technician using a BD LSRFortessa (panel A) or FACSCalibur (panel B) cytometer (Table S2). The FACS data files were re-analyzed in FlowJo version 10 by another single, blinded technician, at the time of this retrospective analysis to ensure consistency in analysis (Fig. S3; Fig. S4). To calculate the circulating number of CD4 T cells, the percentage of CD3+CD4+ cells identified by FACS were multiplied by the total number of circulating lymphocytes quantified by CBC. Similarly, the circulating number of CD8 T cells were calculated by multiplying the percentage of CD3+CD8+ cells by the total number of circulating lymphocytes. The percentages of each of these lymphocyte subpopulations that were CD69+ were generated by the flow cytometry data analysis.

### Data Management and Statistical Analysis

All retrospective macaque data were stored in a custom database built in FileMaker Pro 18 Advanced (Claris International Inc. Santa Clara, CA) specifically for the Retrovirus Laboratory. Microsoft Excel (2001) was used to organize extracted data prior to analysis. A series of mixed effects regression models (MRMs) were conducted in SAS PROC MIXED (version 9.4, Cary, NC) to examine if social housing status influences acute SIV infection (e.g., CD4 T cells, etc.) in juvenile male pigtailed macaques (Table S4). MRMs were used to accommodate repeated laboratory measurements across time that were correlated to different degrees. Additionally, these models account for the fact that different animals were nested in different studies. The full SAS code utilized for this analysis can be made available upon request. Significance was set at *p*<0.05 for a priori analyses, and for exploratory analyses a false discovery rate (FDR) correction was used to control for multiple comparisons. Comparison of the aggregated standard deviation values for all variables included in this analysis (Fig. 3) was done using a two-tailed Wilcoxon matched-pairs signed rank test, with significance set at *p<0.03*. Graphs were constructed using GraphPad Prism (version 8.3.0 for Windows, GraphPad Software, California USA, www.graphpad.com). All data, including deidentified raw data on all parameters, will be available upon request to the corresponding author.

## Acknowledgments

The authors would like to thank members of the Retrovirus Laboratory and Research Animal Resources at Johns Hopkins University, past and present, for their support, especially Samuel Brill, Alisa McNamara, Claire Lyons, Nadine Forbes, Rock Scarborough, Natalie Green, Coffy Bennis, Sara Flemming, and Eric Hutchinson. Thanks for generous ART donation from Abbvie (Abbott), Bristol-Myers Squibb, Gilead, Merck, Jannsen, Roche and ViiV Healthcare. Funding: Grants for Laboratory Animal Science (GLAS), NIH P30 AI094189, P01 AI131306, K01 OD018244, RR00116, P40 OD013117 / U42 OD013117, R01 NS089482, NS097221, NS055651, MH61189, MH070306, NS36911, & RR019995, BSi, Blaustein Pain Foundation.

## Author Contributions

SMGM and KAMP designed this retrospective study, planned, and oversaw all analyses and wrote the manuscript. LHR planned analyses, generated linear mixed model and performed statistics. SMGM, LHR, ENS, SEQ, BWC, RJA, LG, JEC, JLM and KAMP provided input into the experimental design. SMGM, KMM, BWC, ENS and SEQ organized and analyzed data. ENS, SEQ, ML, BB, KAMP, RJA, LG, DRG, MCZ, JEC and JLM performed and analyzed the original studies that generated the data analyzed in this retrospective analysis. LG, DRG, MCZ, JEC and JLM designed the original studies and procured resources for them.

## Materials & Correspondence

Please address correspondence and material requests to Dr. Kelly Metcalf Pate

