## Supplemental Information for "Social Stress Alters Immune Response and Results in Higher Viral Load During Acute SIV Infection in a Pigtailed Macaque Model of HIV"

**Supplementary Tables and Figures:**

| **Controlled Variables** | **Uncontrolled Variables (detailed in Table S2)** |
|---|---|
| Pigtailed macaques | Variety of origins for macaques across all years |
| Juvenile male, 3-5 years old | CBC machine changed in 2015 |
| MANE-A1*084:01 negative | Changes between studies in FACS panel and machine |
| Same room in same facility | Different ART/interventions at d12 PI |
| Same food and water | Unidentified variables associated with pre- & post-2013 |
| Common SIV inoculum stocks | |
| Intravenous route of inoculation | |
| Sampling by same 2 DVMs | |
| Same animal caretaker | |
| Same PCR assay performed by same technician | |
| FACS staining and run by same technician | |

| |
|---|
| FACS re-analyzed by single student |
| Data stored on electronic database |

**Supplemental Table 1. Variability between the singly and socially housed macaque studies**

Controlled and uncontrolled variables that differed between the singly and socially housed macaque studies analyzed in this retrospective study. Differences due to ART regimens used in the different groups were controlled for in the mixed effects regression models. Changes in the CBC machine and in the flow cytometry panels were assessed in post-hoc statistical analyses and found to not impact the outcome of this investigation.

| Database Monkey Group ID | Number of Monkeys | Monkey ID | Year of Inoculation | Age at SIV Inoculation (Years) | Housing Status | Drug Regimen | dPI Treatment Initiation | Source of Monkeys | Hematologic Analyzer | FACs Machine |
|---|---|---|---|---|---|---|---|---|---|---|
| 069 | 4 | 389 | 2015 | 2.9 | Socially | 2.5mg/kg Dolutegravir, 20mg/kg PMPA, 40mg/kg FTC SID SQ | 12 | JHU | Procyte | Fortessa (Panel A) |
| | | 391 | | 2.8 | | | | | | |
| | | 392 | | 2.7 | | | | | | |
| | | 393 | | 2.6 | | | | | | |
| 071 | 4 | 390 | 2016 | 3.6 | Socially | 2.5mg/kg Dolutegravir, 20mg/kg PMPA, 40mg/kg FTC SID SQ, 200mg Maraviroc BID PO | 12 | JHU | Procyte | Fortessa (Panel A) |
| | | 394 | | 3.3 | | | | | | |
| | | 400 | | 3.8 | | | | New Iberia | | |
| | | 401 | | 3.1 | | | | | | |
| 074 | 6 | 409 | 2016 | 3.1 | Socially | 2.5mg/kg Dolutegravir, 20mg/kg PMPA, 40mg/kg FTC SID SQ, 200mg Maraviroc BID PO | 12 | JHU | Procyte | Fortessa (Panel A) |
| | | 410 | | 2.7 | | | | | | |
| | | 411 | | 2.8 | | | | | | |
| | | 412 | | 2.7 | | | | | | |
| | | 413 | | 2.6 | | | | | | |
| | | 414 | | 2.4 | | | | | | |
| 076 | 6 | 422 | 2017 | 2.6 | Socially | 2.5mg/kg Dolutegravir, 20mg/kg PMPA, 40mg/kg FTC SID SQ | 12 | JHU | Procyte | Fortessa (Panel A) |
| | | 423 | | 2.5 | | | | | | |
| | | 424 | | 2.3 | | | | | | |
| | | 425 | | 2.3 | | | | | | |
| | | 426 | | 2.2 | | | | | | |
| | | 427 | | 2.2 | | | | | | |
| 067/073 | 9 | 367 | 2015 | 4.0 | Socially | 30mg/kg PMPA SID SQ for first two weeks then 10mg/kg, 480mg/kg Atazanavir, 10mg/kg Integrase Inhibitor L000870812 BID PO, 24mg/kg Ritonavir BID PO | 12 | JHU | Procyte | Fortessa (Panel A) |
| | | 377 | | 3.8 | | | | | | |
| | | 378 | | 3.6 | | | | | | |
| | | 386 | | 2.8 | | 30mg/kg PMPA SID SQ for first two weeks then 10mg/kg, 480mg/kg Atazanavir, 10mg/kg Integrase Inhibitor L000870812 BID PO, 24mg/kg Ritonavir BID PO | 12 | JHU | Procyte | Fortessa (Panel A) |
| | | 387 | | 2.7 | | | | | | |
| | | 388 | | 2.4 | | | | | | |

| | | | | | | | | | | |
|---|---|---|---|---|---|---|---|---|---|---|
| | | 406 | 2016 | 5.2 | | 30mg/kg PMPA SID SQ for first two weeks then 20mg/kg, 270mg/kg Atazanavir, 10mg/kg Integrase Inhibitor L000870812 BID PO, 24mg/kg Ritonavir BID PO | 12 | New Iberia | Procyte | Fortessa (Panel A) |
| | | 407 | | 3.7 | | | | | | |
| | | 408 | | 3.5 | | | | | | |
| 065 | 6 | 368 | 2015 | 3.9 | Socially | 480mg/kg Darunavir BID PO, 10mg/kg Integrase Inhibitor L000870812 BID PO, 24mg/kg Ritonavir BID PO, 25/50mg/kg Abacavir BID PO | 12 | JHU | Procyte | Fortessa (Panel A) |
| | | 369 | | 3.8 | | | | | | |
| | | 370 | | 3.1 | | | | | | |
| | | 371 | | 3.4 | | | | | | |
| | | 372 | | 3.0 | | | | | | |
| | | 385 | | 3.0 | | | | | | |
| 064 | 6 | 361 | 2013 | 3.9 | Socially | 30mg/kg PMPA SID SQ for first two weeks then 10mg/kg, 270mg/kg Atazanavir, 10mg/kg Integrase Inhibitor L000870812 BID PO, 24mg/kg Ritonavir BID PO | 12 | JHU | IDEXX | Fortessa (Panel A) |
| | | 362 | | 3.9 | | | | | | |
| | | 363 | | 3.8 | | | | | | |
| | | 364 | | 3.6 | | | | | | |
| | | 365 | | 2.6 | | | | | | |
| | | 366 | | 2.4 | | | | | | |
| 054 | 3 | 329 | 2011 | 2.7 | Singly | 30mg/kg PMPA SID IM, 270mg/kg Atazanavir BID PO, 10mg/kg Integrase Inhibitor L000870812 BID PO | 12 | JHU | IDEXX | Fortessa (Panel A) |
| | | 330 | | 2.5 | | | | | | |
| | | 331 | | 2.4 | | | | | | |
| 058 | 3 | 340 | 2012 | 4.4 | Singly | 25mg/kg Fisetin SID PO | 12 | SNBL | IDEXX | Fortessa (Panel A) |
| | | 341 | | 4.4 | | | | | | |
| | | 342 | | 4.4 | | | | | | |
| 050 | 6 | 266 | 2010 | 4.5 | Singly | 12.5mcg Flucanazole SID PO, 5mg Paroxitine SID PO | 12 | New Iberia | IDEXX | Calibur (Panel B) |
| | | 308 | | 3.3 | | | | | | |
| | | 309 | | 3.2 | | | | | | |
| | | 311 | | 3.1 | | | | | | |
| | | 313 | | 3.4 | | | | JHU | | |
| | | 321 | | 4.1 | | | | | | |
| 049 | 3 | 307 | 2009 | 3.9 | Singly | 30mg/kg PMPA SID IM, 270mg/kg Atazanavir BID PO, 10mg/kg Integrase Inhibitor L000870812 BID PO | 42 | Iberia | IDEXX | Calibur (Panel B) |
| | | 310 | | 3.1 | | | | | | |
| | | 312 | | 4.1 | | | | | | |
| 057 | 3 | 337 | 2012 | 5.6 | Singly | 30mg/kg PMPA SID SQ for first two weeks then 10mg/kg, 480mg/kg Darunavir BID PO, 10mg/kg Integrase Inhibitor L000870812 BID PO, 24mg/kg Ritonavir BID PO | 12 | SNBL | IDEXX | Fortessa (Panel A) |
| | | 338 | | 5.5 | | | | | | |
| | | 339 | | 5.5 | | | | | | |
| 052 | 6 | 203 | 2010 | 9.3 | Singly | Untreated | N/a | Yerkes | IDEXX | Calibur (Panel B) |
| | | 268 | | 6.1 | | | | | | |
| | | 272 | | 4.5 | | | | | | |
| | | 274 | | 6.1 | | | | | | |
| | | 320 | | 3.5 | | | | | | |
| | | 265 | | 4.6 | | | | JHU | | |
| 044 | 3 | 291 | 2008 | 3.0 | Singly | Untreated | N/a | Yerkes | IDEXX | Calibur |

| | | | | | | | | | | |
|---|---|---|---|---|---|---|---|---|---|---|
| | | 292 | | 3.0 | | | | | | (Panel B) |
| | | 294 | | 3.0 | | | | | | |
| 061 | 6 | 348 | 2012 | 5.5 | Singly | Untreated | N/a | SNBL | IDEXX | Fortessa (Panel A) |
| | | 349 | | 5.2 | | | | | | |
| | | 350 | | 5.3 | | | | | | |
| | | 351 | | 5.2 | | | | | | |
| | | 352 | | 5.0 | | | | | | |
| | | 353 | | 5.0 | | | | | | |
| 060 | 2 | 345 | 2012 | 5.9 | Singly | 30mg/kg PMPA SID SQ for first 14 days then 10mg/kg, 270mg/kg Atazanavir BID PO, 10mg/kg Integrase Inhibitor L000870812 BID PO, 24mg/kg Ritonavir BID PO | 12 | SNBL | IDEXX | Fortessa (Panel A) |
| | | 347 | | 5.7 | | | | | | |

**Supplemental Table 2. Breakdown of the individual monkey studies included in this retrospective analysis**

This retrospective analysis included 17 different macaque studies—8 of which were included in the socially housed group and 9 of which were in the singly housed group. The majority of the studies initiated an antiretroviral treatment at day 12 post-inoculation; however, the composition of the treatments were different across studies and is thus listed above. The variation in source of the macaques included in each group and the CBC machine and flow cytometry panels utilized for each group are also noted.

| **Panel A** | | | | | |
|---|---|---|---|---|---|
| *Antibody* | *Clone* | *Supplier* | *Catalogue #* | *RRID* | *Fluorochrome* |
| CD3 | SP34-2 | BD Biosciences | 560770 | AB_1937322 | V500 |
| CD4 | S3.5 | Molecular Probes/Life Technologies | Q10007 | AB_11180600 | Qdot 655 |
| CD4 | OKT4 | BioLegend | 317436 | AB_2563050 | Brilliant Violet 650 |
| CD8a | RPA-T8 | Molecular Probes/Life Technologies | Q10152 (discontinued) | AB_1500499 | Qdot 560 |
| CD8a | RPA-T8 | BioLegend | 301038 | AB_2563213 | Brilliant Violet 570 |
| CD69 | FN50 | BioLegend | 310920 | AB_493667 | Pacific Blue |

| **Panel B** | | | | | |
|---|---|---|---|---|---|
| *Antibody* | *Clone* | *Supplier* | *Catalogue #* | *RRID* | *Fluorochrome* |
| CD3 | SP34-2 | BD Biosciences | 552852 | AB_394493 | PerCP-Cy5.5 |
| CD4 | L200 | BD Biosciences | 550628 | AB_393789 | FITC |
| CD8a | RPA-T8 | BD Biosciences | 555366 | AB_395769 | FITC |
| CD69 | FN50 | BD Biosciences | 560738 | AB_1727510 | PerCP-Cy5.5 |

**Supplemental Table 3. Composition of the flow cytometry panels utilized in this retrospective analysis**

Two different flow cytometry panels were used in the studies included in this retrospective analysis. The antibodies and their clone, manufacturer and fluorochrome are listed for each panel.

| | Pre- & Post-Inoculation Interaction | | | Post-Inoculation & ART Interaction | | |
|---|---|---|---|---|---|---|
| | *Social B(SE)* | *Single B(SE)* | *Interaction p-value* | *Social B(SE)* | *Single B(SE)* | *Interaction p-value* |
| Lymphocytes | 1.79 (0.126) | 2.70 (0.14) | <0.0001 | 1.28 (0.13) | 1.53 (0.19) | 0.028 |
| CD4 | 0.59 (0.05) | 0.87 (0.05) | 0.0001 | 0.58 (0.05) | 0.73 (0.06) | 0.293 |
| CD8 | 0.33 (0.04) | 0.57 (0.05) | 0.0002 | 0.16 (0.05) | 0.17 (0.05) | 0.014 |
| CD4/CD8 | 0.20 (0.06) | 0.09 (0.06) | 0.165 | 0.42 (0.05) | 0.60 (0.05) | 0.012 |
| CD69(CD8) | 1.20 (1.44) | 7.09 (2.41) | 0.003 | 0.81 (1.30) | 2.92 (2.30) | 0.005 |
| CD69(CD4) | 1.76 (2.63) | 24.44 (4.08) | <0.0001 | 0.61 (0.73) | 1.45 (1.20) | <0.0001 |

**Supplemental Table 4. Overall interaction values of social versus single housing between pre- and post-inoculation values and between post-inoculation and post-ART values generated by the linear mixed effects regression model**

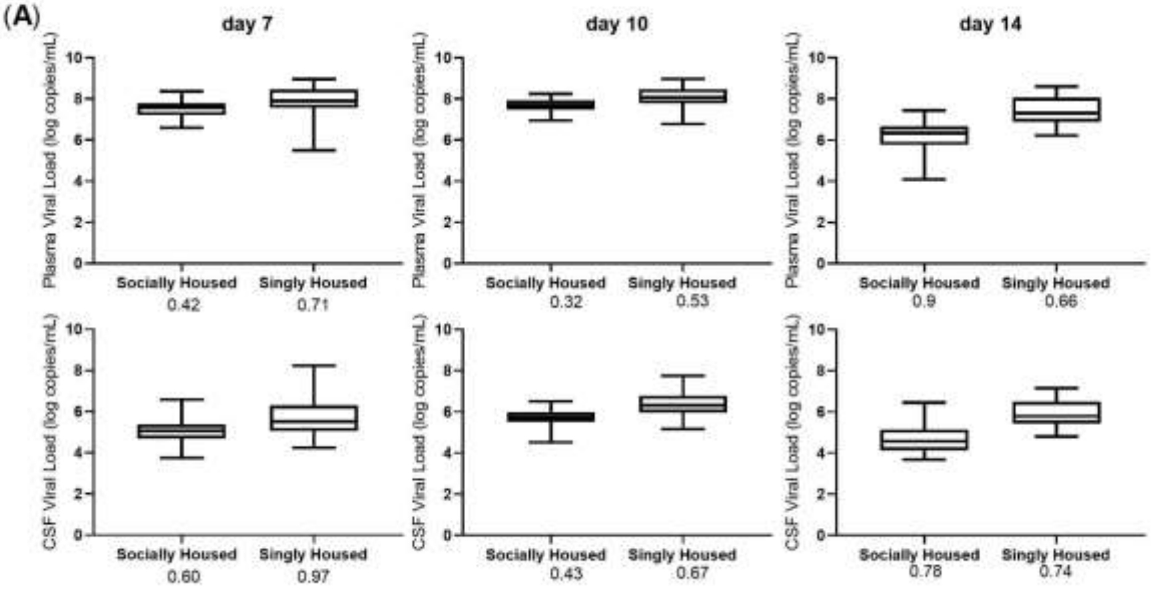

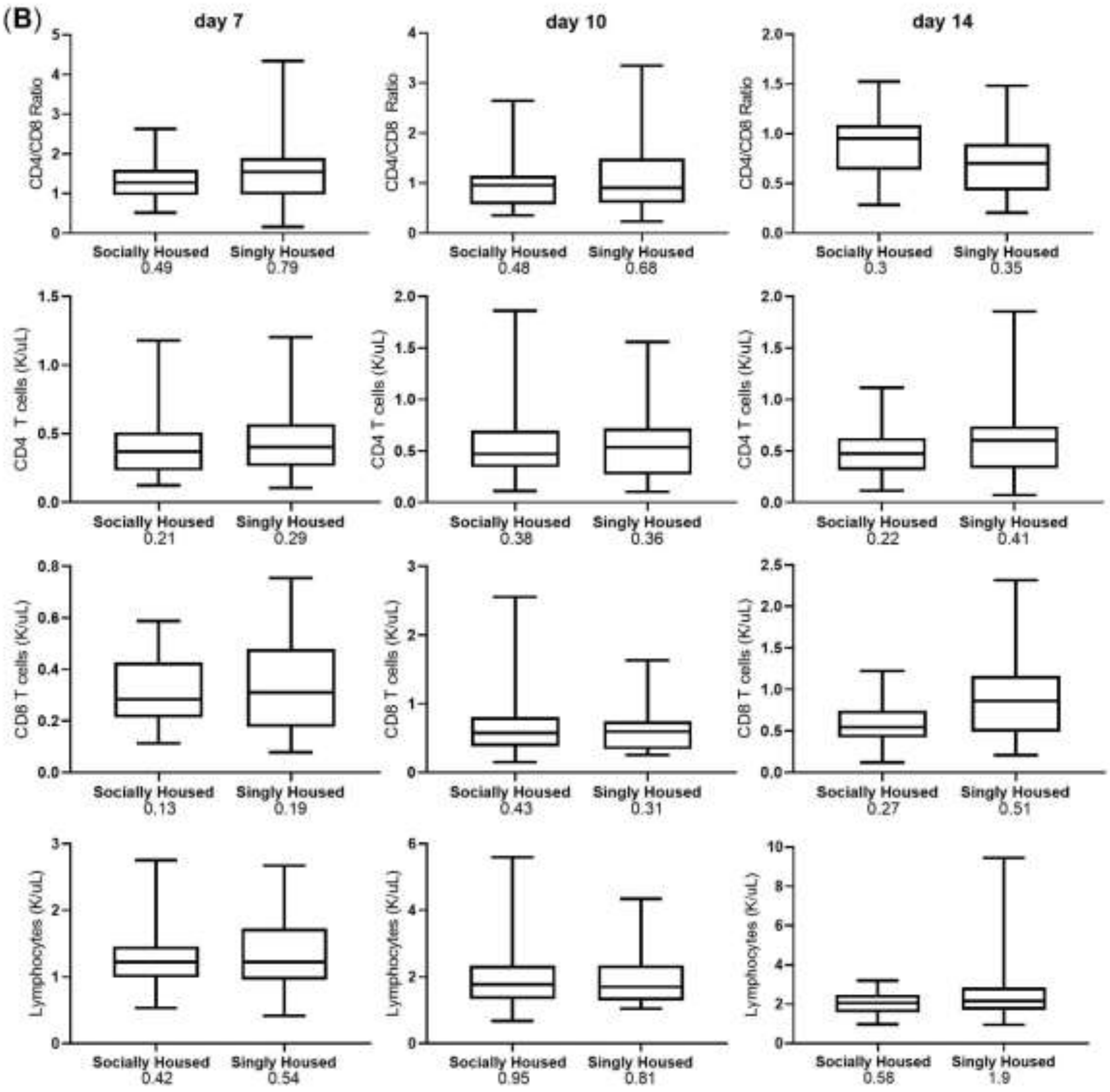
(B)
day 7
day 10
day 14
CD4/CD8 Ratio
CD4 T cells (K/uL)
CD8 T cells (K/uL)
Lymphocytes (K/uL)
Socially Housed
Singly Housed
0.49
0.79
0.48
0.68
0.3
0.35
0.21
0.29
0.38
0.36
0.22
0.41
0.13
0.19
0.43
0.31
0.27
0.51
0.42
0.54
0.95
0.81
0.56
1.9

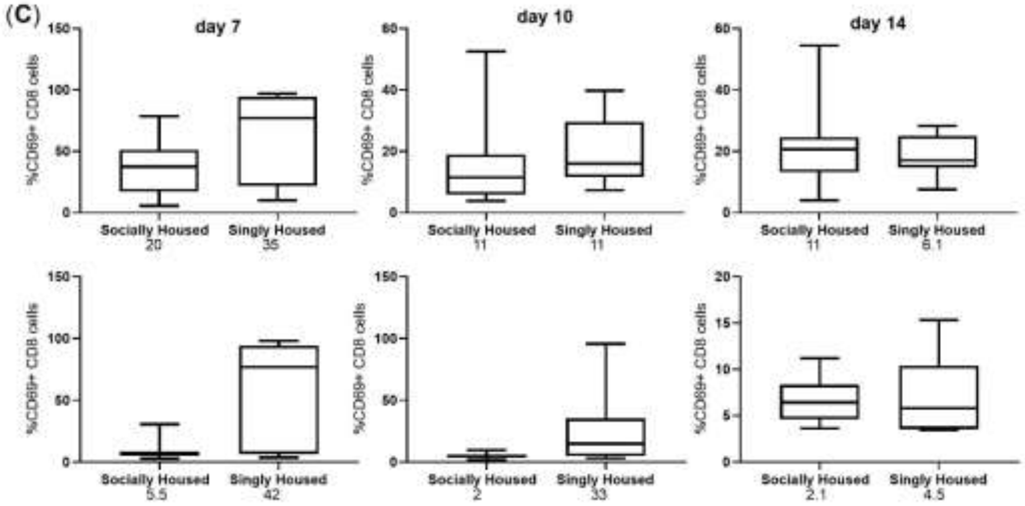

**Supplementary Figure 1. Data variability in acute SIV infection parameters due to social stress**

The standard deviation was calculated for both the singly and socially housed macaque groups at the post-SIV inoculation time points for plasma and CSF viral loads (**A**) CD4/CD8 Ratio, CD4 T cells, CD8 T cells and lymphocytes (**B**) and for the percentage of CD69+ CD8 T cells and for the percentage of CD69+ CD4 T cells. The standard deviation value for each parameter at each time point is listed below the group identification at the x-axis.

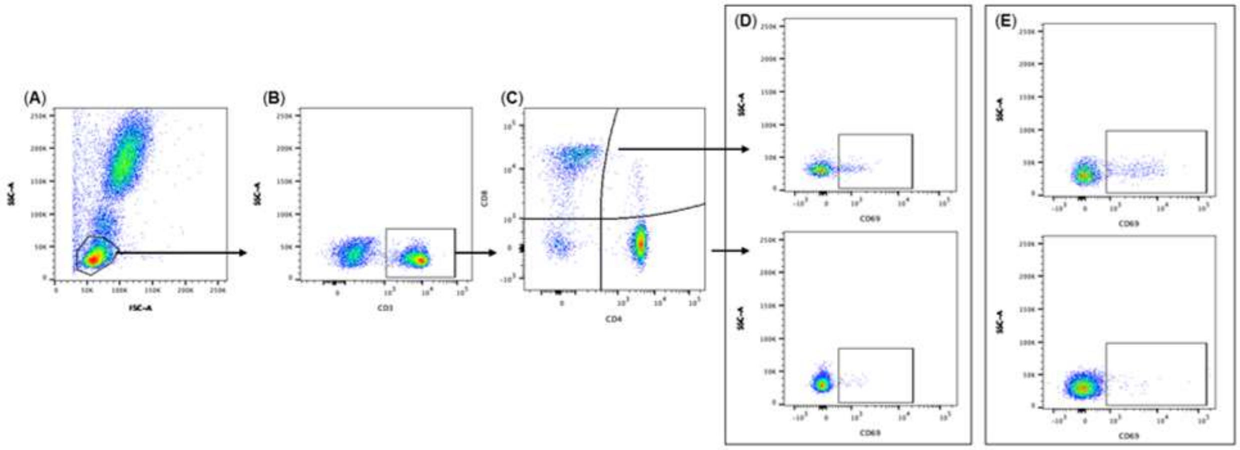

**Supplementary Figure 2. Panel A Flow Cytometry Analysis Gating**

Singlet lymphocytes are gated from whole blood based on forward and side scatter (**A**). CD3+ lymphocytes are identified (**B**) and then gated based on expression of CD4 or CD8 (**C**). Expression of CD69 when either the Pacific Blue fluorochrome (**D**) or the APC fluorochrome is used (**E**) is shown on CD4+ T cells or CD8+ T cells.

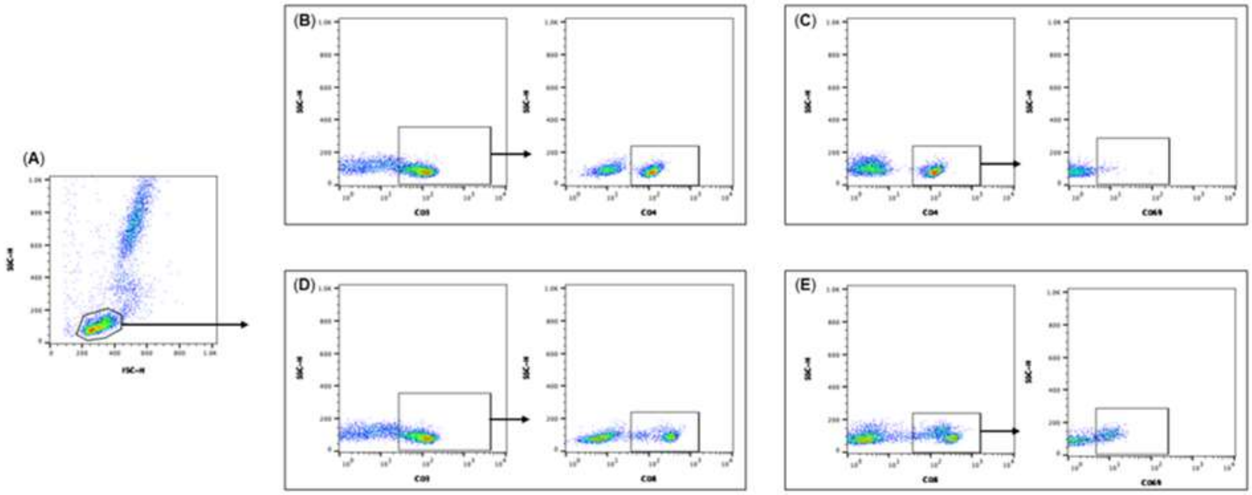

**Supplementary Figure 3. Panel B Flow Cytometry Analysis Gating**

Lymphocytes are gated from whole blood based on forward and side scatter (**A**). For CD4 counts, CD3+ lymphocytes are identified (**B**) and then gated on their expression of CD4. In a separate panel, CD4+ lymphocytes are identified and then gated based on expression of CD69 (**C**). For CD8 counts, CD3+ lymphocytes are identified (**D**) and then gated on their expression of CD8. In a separate panel, CD8+ lymphocytes are identified and then gated based on expression of CD69 (**E**).
